# Impact of an oral amino acid provision on Achilles peritendinous amino acid concentrations in young and older adults

**DOI:** 10.1101/2021.02.12.430945

**Authors:** Chad C. Carroll, Samantha Couture, Dominick O. Farino, Shivam H. Patel, Nathan W.C. Campbell, Julianne Stout, Arman Sabbaghi

## Abstract

Recent studies have indicated that consumption of amino acid-rich compounds can increase tendon collagen content and enhance biomechanical function. Still, it is not clear as to what extent oral consumption of amino acids alters peritendinous amino acid concentrations. Whether aging alters the delivery of amino acids to tendon tissue after oral consumption is also not known. Using microdialysis, we determined the impact of a single oral essential amino acid bolus on Achilles peritendinous amino acid concentrations in younger (n=7; 27±1 yr.) and older adults (n=6; 68±2 yrs.) over four hours. The peritendinous concentration of all amino acids in the beverage except methionine (p=0.136) and glycine (p=0.087) increased with time (p<0.05). Additionally, the concentrations of glycine and arginine were greater in older adults (p≤0.05). We also accessed the impact of amino acid consumption on peritendinous concentrations of pro-collagen Iα1, a marker of collagen synthesis. Pro-collagen Iα1 tended to change with time (p=0.071) but was not altered age (p=0.226). We demonstrate that an oral amino acid bolus leads to modest increases in Achilles peritendinous amino acid concentrations in young and older adults. The concentration of some amino acids was also greater in older adults. However, the amino acid bolus did not significantly impact peritendinous pro-collagen concentrations.

## INTRODUCTION

Recent investigations have implied that tendons are responsive to amino acids. Oral consumption of amino acids may improve clinical outcomes in patients with tendinopathy (15, 30) and optimize tendon adaptations to exercise training (13). Work in rodents has implied that dietary amino acid supplementation can provide an effective means to promote recovery from tendon injury recovery and enhance collagen synthesis (28, 32). In contrast to skeletal muscle, the impact of amino acid consumption on the health of tendon tissue has not been extensively evaluated, especially in humans.

Tendon collagen content is greater in rats given a leucine-rich diet during recovery from malnourishment, suggesting that a diet rich in leucine could increase collagen synthesis. Additionally, a glycine-enriched diet improved tendon collagen content and biomechanical properties in rats after the induction of inflammation via collagenase injection (32). In support of these preclinical findings, patients treated with an arginine-rich oral supplement had modest improvements in pain (15, 25) and rotator cuff repair integrity (15) over those not consuming the supplement. Further, oral supplementation with glycine-rich collagen-peptides improved clinical outcomes in Achilles tendinopathy patients completing a calf-strengthening program (27).

Healthy tendons also appear to be responsive to dietary amino acid/protein-based interventions. Consumption of a vitamin C-enriched gelatin by healthy young men increased serum glycine, proline, and hydroxyproline concentrations, as well as collagen synthesis (30). The serum collected after subjects consumed the vitamin C-enriched gelatin was also used to treat engineered ligaments *ex vivo*. Treated ligaments had greater collagen content and superior biomechanical properties than ligaments treated with serum take before gelatin consumption (30). Consistent with the anabolic effect of leucine on skeletal muscle (10), tendon cross-sectional area (CSA) increased to a greater extent in young adults consuming a leucine-rich whey isolate during a resistance training program compared to a placebo group (13). However, Achilles tendon stiffness was lower in mice given branched-chain amino acids during an exercise intervention than mice on a standard diet (3).

Surprisingly, studies examining the impact of amino acid consumption on tendon properties are limited to young and middle-aged adults. Age-related declines in tendon function (9) and morphological properties (5, 8) and a higher incidence of tendinopathies with age (2, 20) make older adults excellent candidates for amino acid interventions that are proposed to improve tendon properties, yet critical knowledge gaps remain. The extent to which oral consumption of amino acids increases the local delivery of amino acids to tendons is unknown. Knowing this information could guide the development of future oral amino acid beverages to optimize amino acid delivery to tendon tissues. Additionally, whether peritendinous amino acid content is influenced by aging has not been determined. It is well established that oral amino acid provisions increase serum and skeletal muscle amino acid concentrations (17, 22), but such work in human tendon *in vivo* has not been completed.

In this investigation, we utilized microdialysis to determine the impact of oral amino acid consumption on Achilles peritendinous amino acid concentrations in young and older adults (1, 16). We also assessed peritendinous concentrations of pro-collagen Iα1 as a marker of local collagen synthesis in response to the amino acid provision (29). Based on the work described above and work in skeletal muscle (10, 12), we utilized an amino acid bolus rich in leucine and glycine. To our knowledge, no studies have assessed the effects of oral amino acid supplementation on peritendinous amino acid concentrations in young or older adults *in vivo*. To set the framework for future clinical investigations and the optimization of amino acid beverages for therapeutic purposes, establishing the extent to which peritendinous amino acid concentrations change after oral consumption is critical.

## METHODS

### Subjects

Seven young (4 men, 3 women; 27±1 yr.; BMI: 24±1) and six elderly (1 man, 5 women; 68±2 yrs.; BMI: 24±2) adults participated in this research study. Exclusion criteria included body mass index (BMI) greater than 35 kg/m^2^, chronic use of medications known to affect protein or collagen metabolism, a previous history of tendinopathies or diabetes, and sensitivity to study beverage ingredients. All subjects were sedentary (one day or less per week of aerobic or resistance exercise for at least one year) or recreationally active (not training for competitive events). All participants provided voluntary informed written consent. The Institutional Review Board of Purdue University, West Lafayette, IN (IRB#1904022075), approved this research study. This study was registered at ClinicalTrials.gov (NCT04064528).

### Microdialysis

To assess amino acid concentrations in the peritendinous space of the Achilles tendon, we utilized microdialysis (1, 16, 18). For each microdialysis experiment, subjects fasted for 12-hours before arrival at Purdue University. After preparation of the skin with an antiseptic (povidone-iodine) and local anesthetic (lidocaine 1%), an ethylene oxide sterilized microdialysis fiber was inserted in the peritendinous space anterior to the Achilles tendon (1, 16). One hour after fiber insertion, subjects consumed a mixed amino acid beverage prepared from individual amino acid stocks (Ajinomoto Health & Nutrition North America, Inc, Raleigh, NC). Microdialysis samples were collected every 30-minutes for four hours after amino acid consumption.

### Amino Acid Supplement

Subjects received an oral bolus of essential amino acids containing 3.5 grams of leucine (11, 12, 14), 3 grams of proline, 2 g glycine, 1.1 g histidine, 1.0 g isoleucine, 1.55 g lysine, 0.30 g methionine, 1.55 g phenylalanine, 1.45 g threonine, and 1.2 g valine (11). Amino acids (Ajinomoto Health & Nutrition North America, Inc) were mixed in a noncaloric, non-caffeinated carbonated beverage (Crystal Light, Kraft Foods, Inc). The leucine dose was chosen based on work demonstrating its effectiveness at stimulating skeletal muscle protein synthesis (10) and for comparison to previous studies (10, 12). We decided to include greater glycine because of the recent preclinical evidence implying that glycine can improve tendon properties (32). The proline and glycine were meant to provide additional material for collagen synthesis without providing unhealthy levels of these amino acids.

### Sample Analysis

Amino acid concentrations were determined with high-performance liquid chromatography (Agilent Technologies 1100 HPLC System, Santa Clara, CA). Microdialysis samples (15 μl) were deproteinized with 15 μl of 10%TCA and further diluted with 30 μl 0.1N HCl. Diluted samples were immediately centrifugated for 10 minutes at 4°C (10,000 g). The supernatant was removed and transferred to an HPLC vial. Amino acids were eluted using gradient elution with mobile phase A (10 mM Na_2_HPO_4_, 10 mM Na_2_B_4_O_7_, pH 8.2, and 5 mM NaN_2,_ pH 8.2) and mobile phase B (45:45:10 of HPLC-grade acetonitrile, methanol, and water (21). Separation of amino acids was achieved using an Eclipse Plus C18 4.6×100 mm, 3.5μm column (Agilent) with a Restek Ultra C18 Guard Column (Restek Corporation, Bellefonte, PA). Peaks were monitored at 230 nm excitation/450 nm emission (G1321A, Agilent). The concentration of individual amino acid concentrations was determined by comparison with a standard curve (AAS18, MilliporeSigma, St. Louis, MO).

The concentration of pro-collagen 1α1 concentration in the peritendinous space was determined at select time points after amino acid consumption using a DuoSet® ELISA from R&D Systems (DY6220-05, Minneapolis, MN). Due to the large dialysate volume needed for the pro-collagen assay, we could not include every sampling point. Microdialysis samples were diluted 1:9.5 with Reagent Dilutant and assayed in duplicate per the manufacture instructions.

### Statistics

Several participants required a restroom break during the microdialysis experiment resulting in a small number of missed collection points after amino acid consumption. Thus, individual amino acid concentrations were evaluated with a mixed-effects model. For amino acids and pro-collagen, noted residuals were not normally distributed and assumptions of excepted non-constant variance were not correct, thus raw data was log-transformed before analysis. Multiple comparisons were performed using a Fisher’s LSD test. The area under the curve values were compared using an unpaired, two-tailed t-test. All statistical analysis was completed in GraphPad Prism 9.0.1.

## RESULTS

Mean values for all measured amino acids are provided in Table 1. When comparing individual amino acids, peritendinous glycine concentrations (p=0.015, Figure 1) were greater in older adults than young adults after beverage consumption through 150 minutes post-beverage consumption. The age-related difference in glycine concentrations was no longer significant beginning at 180 minutes post-beverage consumption (Figure 1). In contract, glycine did not significantly change with time (p=0.087, Figure 1). Threonine, valine, isoleucine, lysine, leucine, histidine, and phenylalanine concentrations increased with time (p≤0.05, Figure 1) with all of these amino acids returning to baseline by 180 minutes post-beverage consumption. For threonine, valine, isoleucine, lysine, leucine, histidine, and phenylalanine, no differences between young and older adults were detected (p>0.05, Figure 1). The dose of methionine in the study bolus was not sufficient to increase peritendinous concentrations (p=0.136, Figure 1) and no significant difference was noted between young and older adults (p=0.077, Figure 1). No differences across time were noted for the amino acids that were not included in the essential amino acid bolus (p>0.05, Table 1). Additionally, except for arginine (p=0.05), no differences between young and older adults were noted in the peritendinous concentrations of amino acids not included in the essential amino acid bolus.

**Table 1:** Bolded amino acids were included in the study beverage. Values are expressed in μM with mean±SE. Y = young adults; O = older adults. Numbers in first row represent minutes post-amino acid consumption. *p≤0.05 young versus old, ^#^p≤0.05 different from 30 minutes. ^$^p<0.05, main effect for time.

|  | 30 |  | 60 |  | 90 |  | 120 |  | 150 |  | 180 |  | 210 |  | 240 |  |
| --- | --- | --- | --- | --- | --- | --- | --- | --- | --- | --- | --- | --- | --- | --- | --- | --- |
|  | Y | O | Y | O | Y | O | Y | O | Y | O | Y | O | Y | O | Y | O |
| *Glu* | 12±2 | 31±15 | 11±2 | 31±11 | 9±2 | 22±5 | 11±1 | 21±3 | 8±2 | 27±9 | 8±2 | 17±6 | 10±2 | 16±5 | 12±4 | 15±4 |
| *Asn* | 11±2 | 16±2 | 11±2 | 17±2 | 9±2 | 14±2 | 9±1 | 15±1.5 | 8.5±1 | 16±4 | 9±2 | 9±3 | 10±2 | 10±2 | 10±1 | 10±1 |
| *Ser* | 28±4 | 47±9 | 30±5 | 56±10 | 27±6 | 52±10 | 28±5 | 54±10 | 27±4 | 67±22 | 29±7 | 33±8 | 31±6 | 34±7 | 30±5 | 30±3 |
| *Gln* | 273±47 | 555±119 | 292±50 | 612±110 | 241±50 | 468±59 | 253±44 | 557±94 | 245±40 | 614±215 | 272±68 | 302±75 | 298±62 | 322±65 | 279±42 | 320±36 |
| ***His***<sup>\$</sup> | 18±3 | 36±11 | 22±3 | 44±13 | 21±4 | 45±9 | 20±1 | 38±3 | 18±3 | 46±11 | 18±3 | 20±6 | 20±2 | 20±4 | 18±2 | 18±3 |
| ***Gly*** | 66±9 | 133±24* | 85±11 | 182±30* | 87±16 | 197±38* | 98±18 | 199±32* | 84±13 | 217±47* | 86±22 | 109±28 | 87±17 | 107±18 | 79±11 | 102±7 |
| ***Thr*** | 32±9 | 42±8 | 43±10 | 58±8 | 42±9.5<sup>#</sup> | 78±17<sup>#</sup> | 41±8<sup>#</sup> | 78±13<sup>#</sup> | 40±8<sup>#</sup> | 85±22<sup>#</sup> | 42±10 | 41±12 | 42±8 | 42±10 | 89±51 | 38±7 |
| *Arg* * | 17±3 | 30±6 | 16±3 | 33±7 | 11±2 | 27±5 | 14±3 | 30±5 | 11±2 | 31±9 | 13±4 | 15±4 | 15.5±4 | 18±4 | 15±3 | 17±3 |
| *Ala* | 62±13 | 117±26 | 67±14 | 125±13 | 51±14 | 116±7 | 63±13 | 125±10 | 53±11 | 128±22 | 57±17 | 84±23 | 65±17 | 84±18 | 58±10 | 78±10 |
| *Tyr* | 13±2 | 19±2 | 14±2 | 23±2 | 12±3 | 21±2 | 11±2 | 22±2 | 11±2 | 23±4 | 11±2 | 15±4 | 12±2 | 15±3 | 12±2 | 14±1 |
| *Cys* | 70±10 | 127±31 | 92±13 | 166±42 | 88±20 | 159±23 | 105±20 | 177±32 | 82±15 | 202±63 | 85±20 | 116±36 | 94±19 | 118±32 | 86±11 | 105±24 |
| ***Val*** | 61±17 | 65±14 | 79±19<sup>#</sup> | 113±20<sup>#</sup> | 71±15<sup>#</sup> | 104±14<sup>#</sup> | 65±7 | 125±27 | 65±12<sup>#</sup> | 136±49<sup>#</sup> | 64±13 | 56±15 | 65±8 | 56±10 | 62±8 | 52±5 |
| ***Met*** | 10±2 | 14±2 | 12±2 | 19±3 | 10±2 | 18±2 | 10±1 | 17±2 | 9±2 | 18±3 | 9±2 | 12±0.6 | 9±1 | 12±0.9 | 10±2 | 10±0.4 |
| *Trp* | 5±1 | 7±1 | 5±1 | 7±1 | 6±1 | 5±2 | 4±1 | 5±1 | 4±1 | 6±2 | 4±0.9 | 3±0.9 | 6±2 | 3±0.5 | 4±0.9 | 2±0.3 |
| ***Phe*** | 14±3 | 18±3 | 19±3<sup>#</sup> | 30±4<sup>#</sup> | 20±4<sup>#</sup> | 33±4<sup>#</sup> | 18±2<sup>#</sup> | 34±4<sup>#</sup> | 18±3<sup>#</sup> | 38±10<sup>#</sup> | 18±4 | 19±5 | 18±3 | 18±4 | 18±3 | 16±2 |
| ***Iso*** | 16±3 | 18±4 | 26±4<sup>#</sup> | 43±8<sup>#</sup> | 24±5<sup>#</sup> | 42±6<sup>#</sup> | 23±3<sup>#</sup> | 46±10<sup>#</sup> | 19±4<sup>#</sup> | 43±16<sup>#</sup> | 35±17 | 16±5 | 20±3 | 15±3 | 18±4 | 13±2 |
| ***Leu*** | 35±6 | 37±7 | 115±55<sup>#</sup> | 119±26<sup>#</sup> | 74±15<sup>#</sup> | 134±18<sup>#</sup> | 73±10<sup>#</sup> | 143±29<sup>#</sup> | 63±10<sup>#</sup> | 137±46<sup>#</sup> | 57±12 | 54±17 | 55±7<sup>#</sup> | 49±11<sup>#</sup> | 51±7<sup>#</sup> | 44±6<sup>#</sup> |
| ***Lys***<sup>\$</sup> | 32±6 | 45±8 | 41±8 | 67±15 | 36±7 | 59±13 | 34±6 | 64±12 | 33±5 | 61±14 | 33±8 | 31±9 | 33±6 | 31±8 | 30±4 | 29±4 |

**Figure 1:**
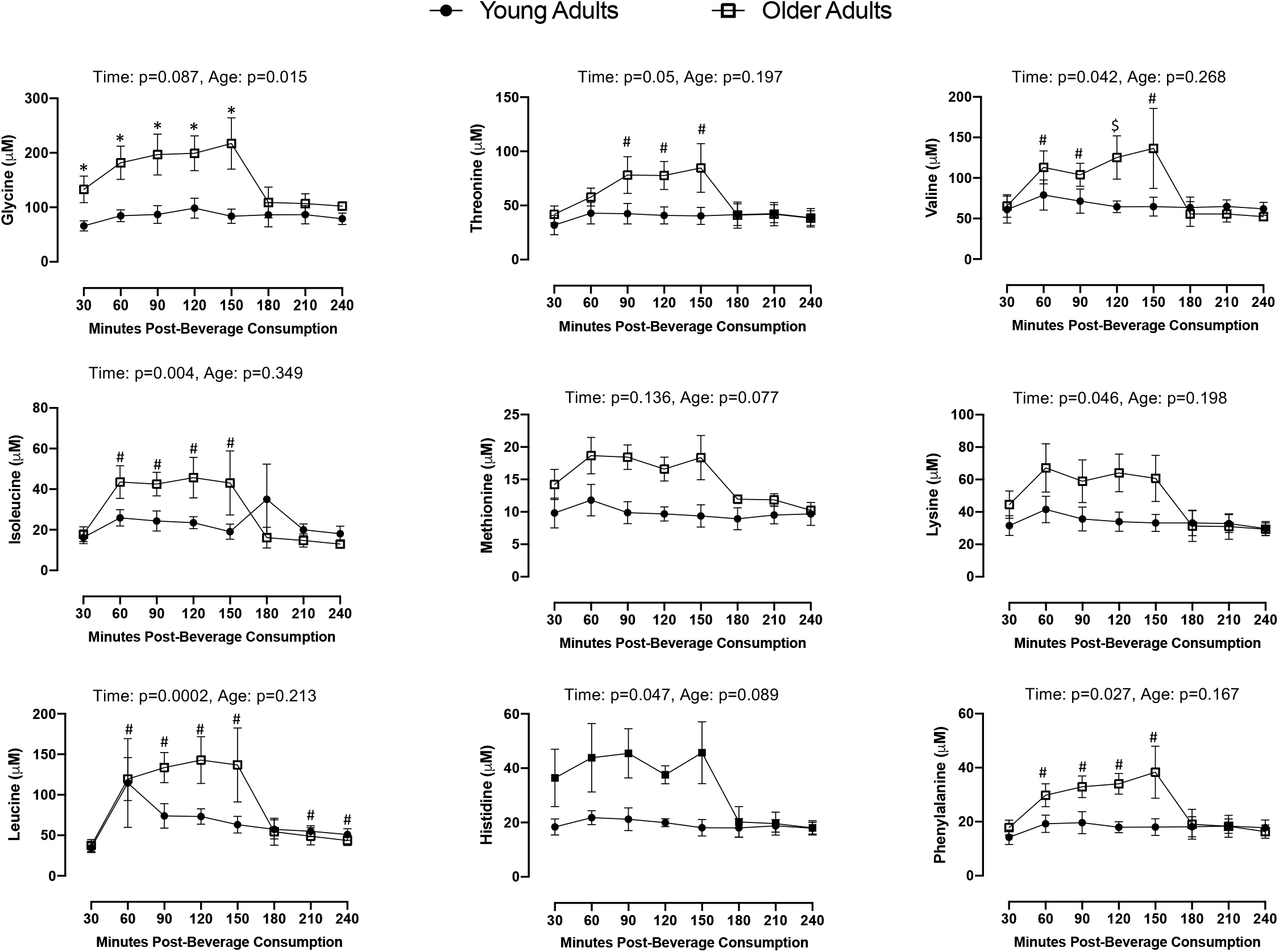
Line graphs of Achilles peritendinous amino acid concentrations over time. Data expressed as mean±standard error. No interactions were noted. Main effect p values for time and age are presented above each figure insert. Multiple comparisons were performed on significant main effects. *p<0.05 versus Young vs. Old; ^#^p<0.05 versus 30 minutes. ^$^p=0.06 versus 30 minutes.

In contrast to the mixed model analysis, AUC values for glycine, histidine, methionine, and phenylalanine were greater in older adults compared to young (Table 2). AUC for lysine and threonine were also higher in older adults but did not reach statistical significance (p=0.06 and p=0.07, respectively). Peritendinous pro-collagen Iα1 concentrations tended to change with time (p=0.071) but were not altered by age (mixed-effect analysis, p=0.226 and AUC, p=0.109, Figure 2).

**Table 2:**

| Amino Acid | Young Adults | Older Adults |
| --- | --- | --- |
| Glycine | 17974±2249 | 33843±3655* |
| Histidine | 4079±414 | 7188±970* |
| Isoleucine | 4948±1086 | 6629±970 |
| Leucine | 14412±3158 | 20294±2889 |
| Lysine | 7234±938 | 10504±1327<sup>#</sup> |
| Methionine | 2069±253 | 3244±239* |
| Phenylalanine | 3821±450 | 5687±603* |
| Threonine | 8574±1267 | 12645±1587<sup>&</sup> |
| Valine | 14092±1831 | 19471±2826 |

Data presented as mean±standard error. \*p<0.05, <sup>#</sup>p=0.0642, <sup>&</sup>p=0.0673

**Figure 2:**
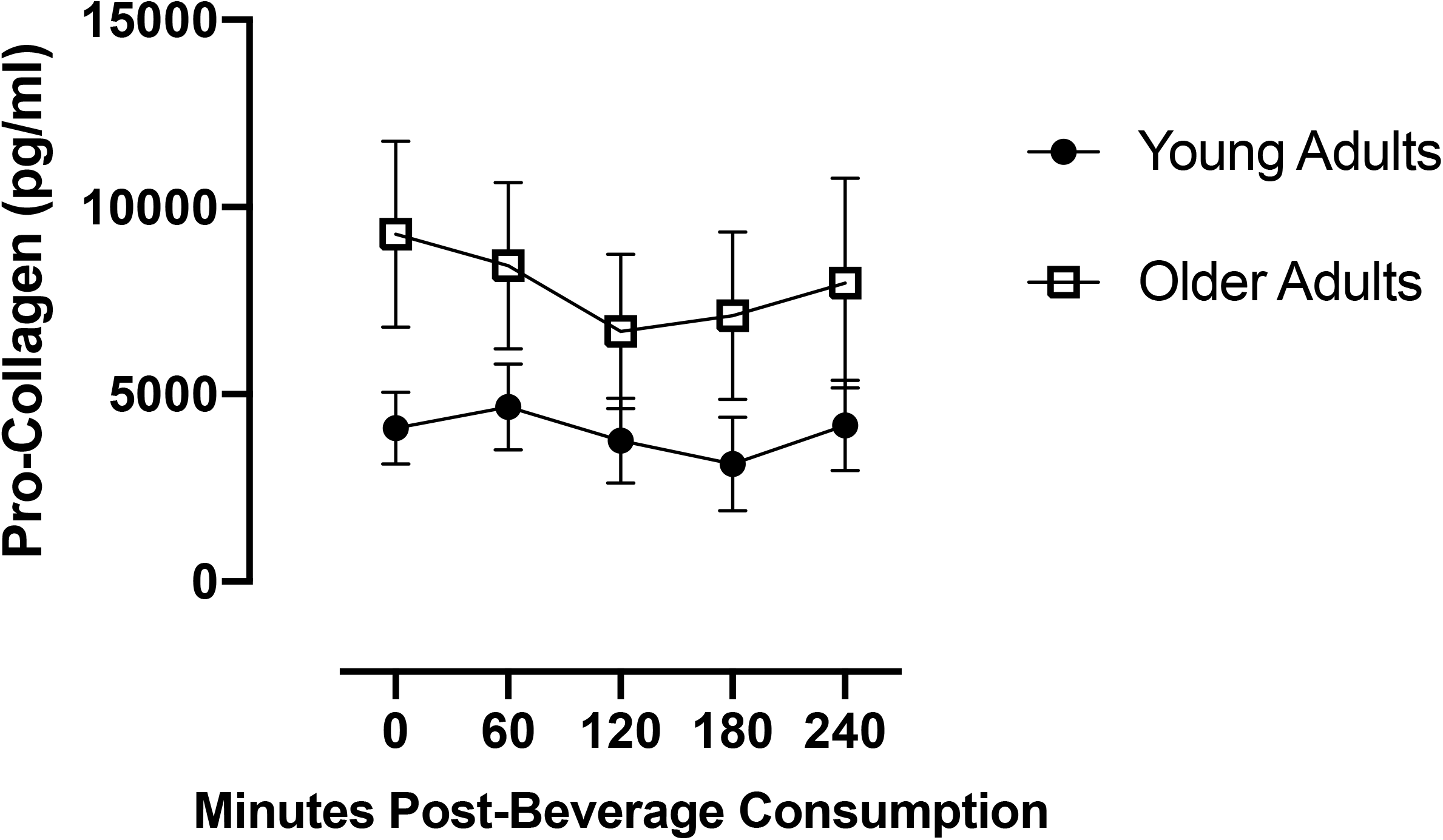
Line graph of Achilles peritendinous pro-collagen concentrations over time. Data expressed as mean±standard error.

## DISCUSSION

It is well-established that an elevation of serum amino acids can have a significant anabolic effect on skeletal muscle (6, 11, 31). A limited number of investigations in humans and rodents have indicated that tendon tissue is also sensitive to elevations in amino acids. Amino acid or protein beverages have increased tendon CSA in adults completing resistance training (13) and improved outcomes during tendinopathy rehabilitation (15, 30). In rodents, amino acids solutions, especially those rich in leucine or glycine, improve tendon properties in various disease models (28, 32). Consumption of amino acid or protein beverages increases serum and skeletal muscle amino acids. However, to our knowledge, the extent to which oral amino acid ingestion alters local (peritendinous) concentrations of amino acids in humans has not been described. The potential impact of amino acids on tendon properties in older adults is also relevant, given the changes in tendon properties with aging (5). Thus, we also considered the impact of aging on the ability of oral amino acids to increase peritendinous amino acid concentrations. Additionally, we determined if a provision of amino acids would increase the peritendinous concentration of a common collagen synthesis marker.

In this study, we demonstrate that amino acids can be detected in the Achilles peritendinous space of humans. The concentrations of the amino acids not included with the beverage were stable across time, suggesting a minimal impact of fiber insertion (Table 1). Of the amino acids in the drink, all increased with time (p<0.05), except methionine and glycine. The change in Achilles peritendinous amino acid concentrations across time followed a similar pattern to those previously reported in serum and skeletal muscle (1, 10) with levels peaking at 1-2 hours post-ingestion then returning to baseline levels at approximately three hours (Figure 2).

While direct statistical comparisons are not possible, the observed basal peritendinous amino acid concentrations (Table 1) were substantially lower than previously reported in serum and skeletal muscle (7, 10). Additionally, the peak concentrations obtained in the peritendinous space after consuming the amino acid bolus were generally lower than skeletal muscle or serum (10). For example, utilizing a similar amino acid bolus, Dickinson et al. (10) reported peak serum concentrations of approximately 1000 μM and 250 μM for leucine and phenylalanine, respectively. In comparison, the highest concentration obtained in the Achilles peritendinous space for leucine was 115 μM for young and 143 μM for older adults. Phenylalanine Achilles peritendinous concentrations peaked at 20 μM for young and 38 μM for older adults. We did not account for probe recovery due to the large number of analytes examined. However, we have previously reported a probe recovery range of 50-65% for the amino acid sarcosine (16). Even accounting for estimation of recovery, the values obtained in the Achilles peritendinous space are lower than those obtained in serum or skeletal muscle. Further, the increase in peritendinous amino acid concentrations after the oral bolus was modest when compared to serum or skeletal muscle. Even with the larger dose of glycine, we did not observe a significant increase in peritendinous glycine concentrations with time. This suggests that delivery of amino acids to the Achilles tendon is limited when compared to skeletal muscle and serum. Larger oral doses of amino acids may be needed to achieve greater benefits for tendons.

A lower concentration of amino acids after oral consumption is consistent with our previous work with acetaminophen (16). Achilles peritendinous levels of acetaminophen achieved maximum values that were approximately 50% lower than those seen in serum or skeletal muscle after oral consumption of the drug (24). It is not yet clear why the peritendinous concentration of compounds is lower than serum or skeletal muscle compared to the Achilles peritendinous space. While the tendon is poorly vascularized, blood flow in the peritendinous space during non-exercise conditions is similar to the surrounding calf muscle (4) implying that delivery of amino acids to the tendon would not be impaired.

Interestingly, when accounting for the area under the curve, the concentration of several of the amino acid included in the beverage achieved higher peritendinous concentrations in older adults than in young (Figure 1). Arginine, which was not included in the amino acid bolus, was also higher in the older participants compared to young. A greater baseline peritendinous amino acid concentration is surprising as serum amino acid concentrations are typically lower in older adults than young (26).

With the apparent anabolic effect of leucine and glycine on tendon collagen (32) and mass (13), we evaluated peritendinous pro-collagen Iα1 as a marker of collagen synthesis. Even with the large proportion of leucine and glycine in the amino acid beverage, we did not observe a significant increase in peritendinous levels of pro-collagen Iα1 (p=0.071). One limitation of the peritendinous microdialysis approach is that we assume that any pro-collagen detected is a reflection of intratendinous events. Extensive work by Langberg and colleagues (19) has validated the peritendinous microdialysis technique providing evidence that peritendinous measures correlate with intratendinous measures. However, using stable isotope methods to directly assess collagen synthesis (23) would provide more detailed results but would require invasive tissue sampling.

In future work, it would be interesting to determine if a different source of protein (e.g., whey or soy) or variations in beverage amino acid content would alter peritendinous amino acid concentrations, as reported for serum (31). The limited rodent and human work suggest that tendon tissue is indeed sensitive to serum amino acids changes. However, given the lower concentrations obtained in the peritendinous space, we should carefully explore the dose-response impact of amino acids on tendon tissue using *in vitro* or *ex vivo* models. Such information is critical for optimizing oral doses of amino acids for human consumption to maximize the stimulation of collagen synthesis. Also, understanding the anabolic potential of each amino acid would aid in optimizing beverage content. It would be exciting to determine if a larger bolus or repeated smaller doses of amino acids would increase peritendinous amino acid concentrations to a greater extent than seen in the current investigation. Understanding the impact of these beverages on peritendinous amino acid concentrations will optimize beverage content to maximize the clinical benefit of such compounds.

While much work has attempted to define an optimal protein dose to stimulate skeletal muscle protein synthesis, such information is not yet available for the tendon. Developing a nutritional cocktail to optimize tendon and skeletal muscle health is appropriate given the importance of both tissues to overall musculoskeletal function. The small number of human studies suggest the exciting possibility that amino acid beverages could be optimized to increase peritendinous amino acids level for optimization of tendon health and recovery while providing the established benefits for skeletal muscle.

## Acknowledgments

We thank the subjects for their participation and commitment to the investigation. This project was funded by Purdue University Research Initiative Funds to CCC. The study was designed by CCC; data were collected and analyzed by all authors; data interpretation and manuscript preparation were undertaken by CCC and AS. All authors approved the final version of the paper.

